# Long-read sequencing reveals rapid evolution of immunity- and cancer-related genes in bats

**DOI:** 10.1101/2020.09.09.290502

**Authors:** Armin Scheben, Olivia Mendivil Ramos, Melissa Kramer, Sara Goodwin, Sara Oppenheim, Daniel J Becker, Michael C Schatz, Nancy B Simmons, Adam Siepel, W Richard McCombie

## Abstract

Bats are exceptional among mammals for their powered flight, extended lifespans, and robust immune systems. To investigate the genomic underpinnings of unique bat adaptations, we sequenced the genomes of the Jamaican fruit bat (*Artibeus jamaicensis*) and the Mesoamerican mustached bat (*Pteronotus mesoamericanus*) and compared them to a diverse collection of 13 additional bat species together with other mammals. We used the Oxford Nanopore Technologies long-read platform to generate highly complete assemblies (N50: 28-29Mb) and facilitate analysis of complex genomic regions containing duplicated genes. Using gene family size analysis, we found that the type I interferon locus was contracted by eight genes in the most recent common ancestor (MRCA) of bats, shifting the proportion of interferon-ω to interferon-α and making interferon-ω the most common type I interferon in bats. Antiviral genes stimulated by type I interferons were also rapidly evolving, with interferon-induced transmembrane genes experiencing a lineage-specific duplication and strong positive selection in the gene *IFIT2*. Moreover, the lineage of phyllostomid bats showed an unprecedented expansion of *PRDM9*, a recombination-related gene also involved in infection responses, raising the possibility that this gene contributes to bat antiviral defenses. These modifications in the bat innate immune system may be important adaptations allowing them to harbor viruses asymptomatically. We additionally found evidence of positive selection on the branch leading to the MRCA of bats acting on 33 tumor suppressors and six DNA repair genes, which may contribute to the low cancer rates and longevity observed across bats. These new genomic resources enable insights into the extraordinary adaptations of bats, with implications for mammalian evolutionary studies and public health.

## Introduction

Bats (order Chiroptera) form the second largest order of mammals and are known for a wide variety of remarkable adaptations including powered flight^1^, laryngeal echolocation^2,3^, unusual longevity^4^, and low rates of cancer^5^. Bats are also hosts of diverse viruses^6,7^ and have played roles in outbreaks of emerging zoonotic viruses including Marburg virus^8^, Nipah virus^9^, and both severe acute respiratory syndrome coronavirus 1 (SARS-CoV-1)^10^ and SARS-CoV-2^11^, either through direct human contact or via bridge hosts. It has been suggested that their tolerance of many viral infections stems from unusual features of their innate immune response^12^. Together, these adaptations make bats a powerful system for investigating a wide variety of genotype-to-phenotype relationships, including several with implications for human health. For example, by better understanding the mechanisms of the bat immune system that allow them to tolerate viral infections^5^, we may be better able to prevent zoonotic outbreaks. In addition, comparative genomic analyses of bats and cancer-susceptible mammals may shed new light on the causes of cancer and links between cancer and immunity^13^. Importantly, such studies of bats and other non-model organisms are highly complementary to studies based on mouse models, which are far more amenable to experimental manipulation but exhibit fewer natural adaptations relevant to human disease.

With these goals in mind, investigators have sequenced and assembled the genomes of at least 43 bat species over the past decade (**Table S1**). Recently, sequencing efforts in bats have been accelerated by the Bat1K global genome consortium^14^ (http://bat1k.com), DNA Zoo^15^ (https://www.dnazoo.org/) and Vertebrate Genome Project (https://vertebrategenomesproject.org). These new genome sequences have revealed numerous intriguing features of the immune systems of bats^12,16–22^. In particular, several genes with key roles in the innate immune system appear to have adaptively evolved in bats, including primary lines of inducible host defences such as pathogen sensors^16,23^, type I interferons (IFNs)^12,24^ and antiviral genes^25^. Specifically, bats have lost the mammalian PYHIN DNA-sensing gene family^17,26^, they show evidence of positive selection in pathogen-sensing Toll-like receptors (TLRs) ^23^, and they display copy-number variation in type I IFN cytokines^12,24^, which are induced by TLRs. Bat-specific modifications in tumor suppressors, DNA damage checkpoint-DNA repair pathway genes^17^, and growth hormone^27^ may be associated with cancer resistance. It is thought that these adaptations in innate immunity and cancer resistance may have arisen as a result of coevolution of bats with viruses^18,28^, and that a need for enhanced DNA repair in the face of elevated reactive oxygen species (ROS) may have been a consequence of powered flight^17^.

In this study, we augment previously existing genome sequences with new Oxford Nanopore Technologies (ONT)-based long-read assemblies for the Jamaican fruit bat (*Artibeus jamaicensis*) and the Mesoamerican mustached bat (*Pteronotus mesoamericanus*) (**Figure S1**). We present a comprehensive analysis of these genome sequences together with 13 previously assembled bats and other mammalian genomes. Importantly, long-read assemblies enable dramatic improvements in the characterization of gene duplications and losses, and of genomic repeats^29–31^. These benefits are of particular value in studies of mammalian immunity-related genes, many of which fall in highly repetitive genomic regions including large arrays of duplicated genes^32^. Our comparative genomic analysis of these genome sequences provides several new insights into unique features of innate immune response and cancer resistance in bats.

## Results

### Genomic structure of *A. jamaicensis* and *P. mesoamericanus*

Except for two recently published studies^12,18^, most previous genomic investigations in bats have relied on short-read assemblies based on Illumina DNA sequencing technology, which has limited the study of complex regions. In this case, we were able to leverage the ONT long-read sequencing platform and an optimized flye^33^-PEPPER^34^-POLCA^35^ assembly and polishing strategy (**Methods**) to generate reference-quality assemblies for *A. jamaicensis* and *P. mesoamericanus* with contig N50 values of 28-29Mb (**Figure S2**, **Table S2**) and POLCA consensus accuracy >99.99%. Using EVidenceModeler, we annotated 21,621 genes in *A. jamaicensis* and 21,269 genes in *P. mesoamericanus*. Based on a BUSCO protein assessment of our annotations, the gene sets in both bats are highly complete at 98.3% and 98.2% respectively, comparable to the values of 97.4-98.3% reported for six recent PacBio-based bat assemblies (**Figure S3**). Notably, all of these long-read bat assemblies have BUSCO scores approaching those of the human (99.9%) and mouse (99.9%) annotations. Orthofinder orthology detection produced 19,935 orthogroups for 15 bats and 5 outgroup mammals, of which 12,517 single-copy orthogroups were set aside for our positive selection analysis (below). Total fractions of 39.2% and 37.9% of the *A. jamaicensis* and *P. mesoamericanus* genomes consisted of repeats, respectively, with 0.4% in each genome attributed to recently active transposons including hAT, TcMariner, and piggyBac elements (**Figure S4**, **Table S3**). We also detected non-retroviral endogenous viral elements, predominantly derived from Bornaviridae and Parvoviridae (**Table S4**). We provide our annotations, aligned evidence, and multiple genome alignments as a public UCSC genome browser instance (http://compgen.cshl.edu/bat/).

### Gene family expansion and contraction analysis

Changes in gene family size have played an important role in shaping the immune systems of bats^17,26^. To facilitate further analysis of gene family expansions and contractions, we focused on our new ONT-based assemblies and the previously published long-read bat genome sequences. By comparing these bat genomes with mammalian outgroups (human, mouse, dog, pig, and horse), we identified four expanded gene families and 47 contracted gene families in the most recent common ancestor of bats (hereafter, the “bat MRCA”; **Table S5**). Twenty-five of the gene families changing size on the bat ancestral branch were related to immune system processes, including the previously identified PYHIN gene family (PTHR12200)^26^, which we confirmed to be entirely absent in all bats including the two newly sequenced species.

### The type I interferon locus is contracted in bats

The type I IFN immune response is a critical component of the mammalian innate immune system, and is responsible for activating the expression of a battery of antiviral genes following induction by pathogen-sensing components such as PYHINs, TLRs, and *cGAS*-*STING*^36^. Previous comparative analyses of the type I IFN locus have shown that it is highly structurally variable in bats and other mammals^24^, with some bats, such as *Pteropus alecto,* showing a contraction^24^, whereas others such as *Rousettus aegyptiacus*^12^, *Pteropus vampyrus,* and *Myotis lucifugus*^37^ show evidence of expansions. However, this locus is generally large and highly duplicated across mammals (e.g., in humans, it spans ∼400kb and contains 16 IFN genes, including 13 IFN-α genes and one IFN-ω gene), making it challenging to assemble and analyze.

Using our expanded set of long-read-based bat genomes, we found clear evidence of a major contraction (-8 genes; Viterbi method, see ref.^38^, *p*=6.4e-11) of the type I IFN gene family in the bat MRCA (**Figure 1**, **Figure S5**). Interestingly, we found that this contraction was driven specifically by loss of IFN-α genes, with gene counts of 0-4 in bats compared to 10-18 in the outgroup mammals (**Figure 1**). In contrast, IFN-ω gene counts in the bat ancestor were largely unchanged (Viterbi method, *p*=0.51), ranging from 3-7 in bats and 0-8 in other mammals. As a consequence, IFN-ω was 11-fold enriched relative to IFN-α in bats compared to the other mammals (Fisher’s exact test, *p*=4.5e-11, odds ratio=10.80) and this enrichment was observed in every bat species (**Table S6**). Considering the relative ligand-binding and antiproliferative properties of IFN-ω and IFN-α^39,40^, these changes in gene number could potentially be responsible for more potent responses to viral infections in bats relative to other mammalian orders (see **Discussion**).

**Figure 1.**
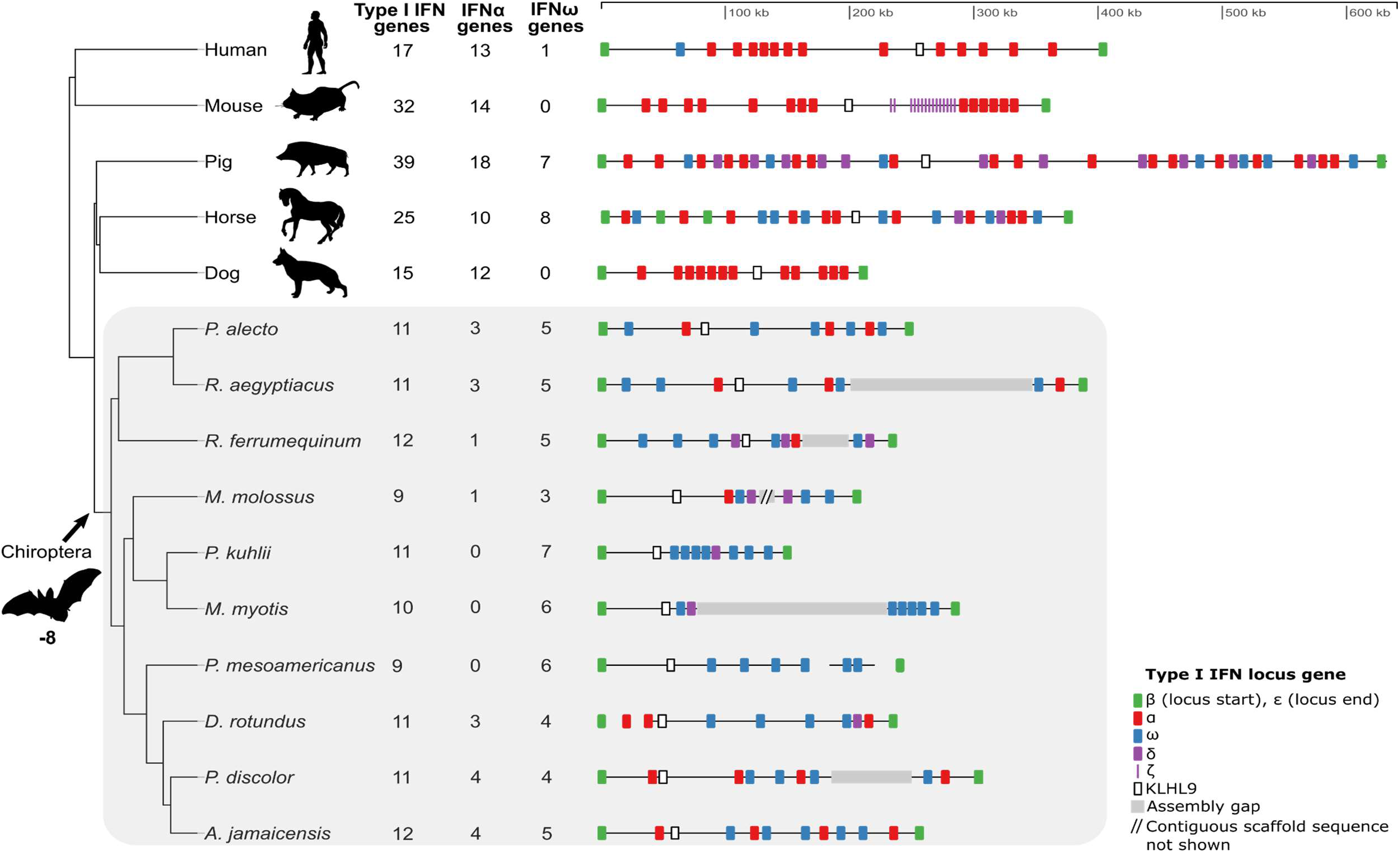
Contraction of the type I interferon (IFN) locus in bats compared to other mammals. A loss of eight genes in the bat MRCA was estimated by CAFE^38^ (Viterbi method, *p*=6.4e-11). The reduction in locus size occurred together with a significant loss of IFN-α genes but not IFN-ω genes in the bat MRCA (Fisher’s exact test, *p*=4.5e-11, odds ratio=10.80). The type I IFN loci in bats (grey background) are shown for long-read based assemblies as well as a BAC-based locus assembly^24^ (*P. alecto*) and an Illumina-based assembly^19^ (*Desmodus rotundus*). Abbreviated species names: *Rhinolophus ferrumequinum*, *Molossus molossus*, *Pipistrellus kuhlii*, *Myotis myotis*, and *Phyllostomus discolor*.

### Antiviral interferon-induced transmembrane genes are expanded in yangochiropteran bats

An important downstream consequence of the activation of type I IFNs is the expression of various antiviral IFN-stimulated genes (ISGs)^41^. In bats, several of these genes, such as *tetherin* and *APOBEC3,* have also been shown to be under positive selection^42^ or are duplicated^18^. Among antiviral ISGs, we observed an expansion of the immune-related IFN-induced transmembrane (IR-IFITM) gene family (+2 genes; Viterbi method, *p*=1.6e-3) on the branch leading to the Yangochiroptera, the suborder that includes most microbats (**Figure 2**). The IR-IFITMs—which have previously been reported to be under positive selection in bats^43^—are potent broad-spectrum antiviral factors that help to prevent infection before a virus passes the lipid bilayer of the cell^44,45^. By applying a dN/dS-based branch-site likelihood ratio test (**Methods**), we confirmed using our data that the IR-IFITMs show evidence of positive selection in the bat MRCA (*p*=6.4e-3). Furthermore, we identified seven particular codon sites that show signs of episodic diversifying selection (**Table S7**), including three sites in the CD225 domain (codons 68, 70, and 117). Notably, codons 68 and 70 in the CD255 domain are among several sites previously shown to be critical for blocking viral infection^43^. Furthermore, the methionine-to-phenylalanine substitution at codon 68, occurs in an amphipathic helix previously shown to be essential for blocking viral infection^46^. Together these observations of gene duplications and positive selection in functional domains in IR-IFITMs suggests that this gene family may have played an important role in the evolution of antiviral responses in bats.

**Figure 2.**
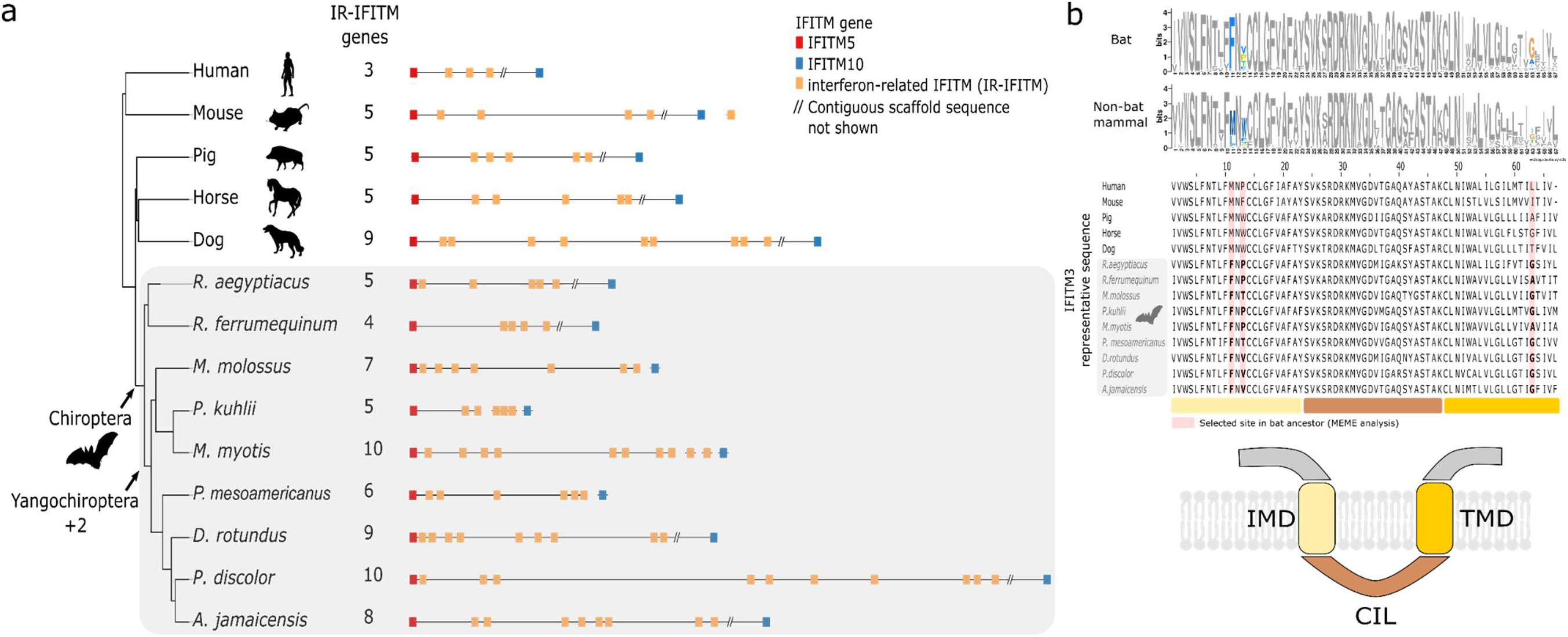
IFITM gene family expansion and positive selection associated with the bat antiviral immune response. a) Phylogeny of bats and other mammals, showing a significant increase in gene copy number at the IFITM locus in yangochiropteran bats based on CAFE^38^ analysis. Immune-related interferon-induced transmembrane (IR-IFITM) genes are shown in yellow. b) Three codon sites in the IFITM transmembrane domains IMD and TMD in bats show evidence of positive selection (see also **Table S7**). The sequence logo (top) compares a 67-amino-acid region spanning these domains with orthologous regions from other mammals (selected sites in color). The sequence alignment (middle) compares human *IFITM3* with the most similar representative IR-IFITM ortholog from the other species (selected sites highlighted with red background and shown in bold). The protein model (bottom) is based on a published transmembrane topology^47^.

### Expansion of *PRDM9* in phyllostomid bats and expansion of heat shock proteins in *P. mesoamericanus*

A third gene family to emerge from our survey of gene expansions and contractions was *PRDM9* (**Table S8**), which specifies the location of meiotic recombination sites^48^ and is known to evolve rapidly in vertebrates^49^. *PRDM9* may play a role in speciation^50^ and is also upregulated upon viral infection^51^. We found that *PRDM9* experienced a striking expansion in phyllostomid bats (+5 genes; Viterbi method, *p*=1.2e-09; **Figure S6**), far beyond anything observed in other mammals. Intriguingly, many of the *PRDM9* copies in phyllostomid bats have lost the KRAB domain (**Table S9**), which is thought to play an important role in recruiting the recombination machinery^49^, suggesting they may have alternate functions. Finally, we observed a major expansion in *P. mesoamericanus* of heat-shock proteins across multiple gene families, including heat-shock protein 70kDa (PTHR19375, +11 genes; Viterbi method, *p*=8.2E-14). Interestingly, this expansion was largely restricted to *P. mesoamericanus* and was not observed across other bats. Overexpression of heat-shock proteins can modulate immune responses^52^, therefore this duplication may have implications for immunity in *P. mesoamericanus*.

### Positive selection analysis

Having identified signatures of positive selection at the amino-acid level in several gene families of interest, we applied similar branch-site likelihood ratio tests^53–55^ genome-wide, focusing on 12,517 single-copy orthologs present in bats and outgroup mammals. Because we were most interested in adaptations shared by most bats, we mainly focused on a test for positive selection on the branch leading to the bat MRCA, where we expected to have reasonably good statistical power. However, we also tested for positive selection on the lineages leading to each of the two newly sequenced bats. Observing a highly skewed distribution of nominal *p*-values (*p*>0.98 in 86% of tests), we opted to follow a recent study^56^ and omit a correction for multiple comparisons across orthologs, instead adjusting only for testing on three different branches (but see **Table S10** and **Data S1** for more conservative adjusted *p*-values). Based on this strategy, we identified 468 positively selected genes (PSGs) with adjusted *p*<0.05 in the bat MRCA (**Figure 3, Table S10**). This number is roughly comparable to the 298 PSGs recently identified for the Phyllostomidae bat lineage using a similar filtering approach^56^ but somewhat larger than the numbers found on the branch leading to the bat MRCA in recent studies that used more stringent criteria (181 PSGs^20^ and 23 PSGs^18^). These PSGs were strongly enriched for immune-related functions (**Table S11**), including the major GO biological processes “regulation of inflammatory response” (GO:0050727, *p*=6.1e-4) and “innate immune response” (GO:0045087, *p*=6.2e-4). In total, 125 PSGs (27% of the 468) were annotated with the parent term “immune system process” (GO:0002376). The bat MRCA branch was also enriched for PSGs involved in “positive regulation of reactive oxygen species (ROS)” (GO:2000379, *p*=3.5e-4), possibly suggesting adaptations associated with heightened metabolic rates owing to flight (see **Discussion**). Moreover, we detected six autophagy-related PSGs (**Table S10**), including autophagy regulator *ATG9B* which was previously implicated in bat longevity^57^. Below, we further discuss some specific PSGs falling in two major classes: (1) pathogen sensors, cytokines, and antiviral genes; and (2) DNA repair genes and tumor suppressors.

**Figure 3.**
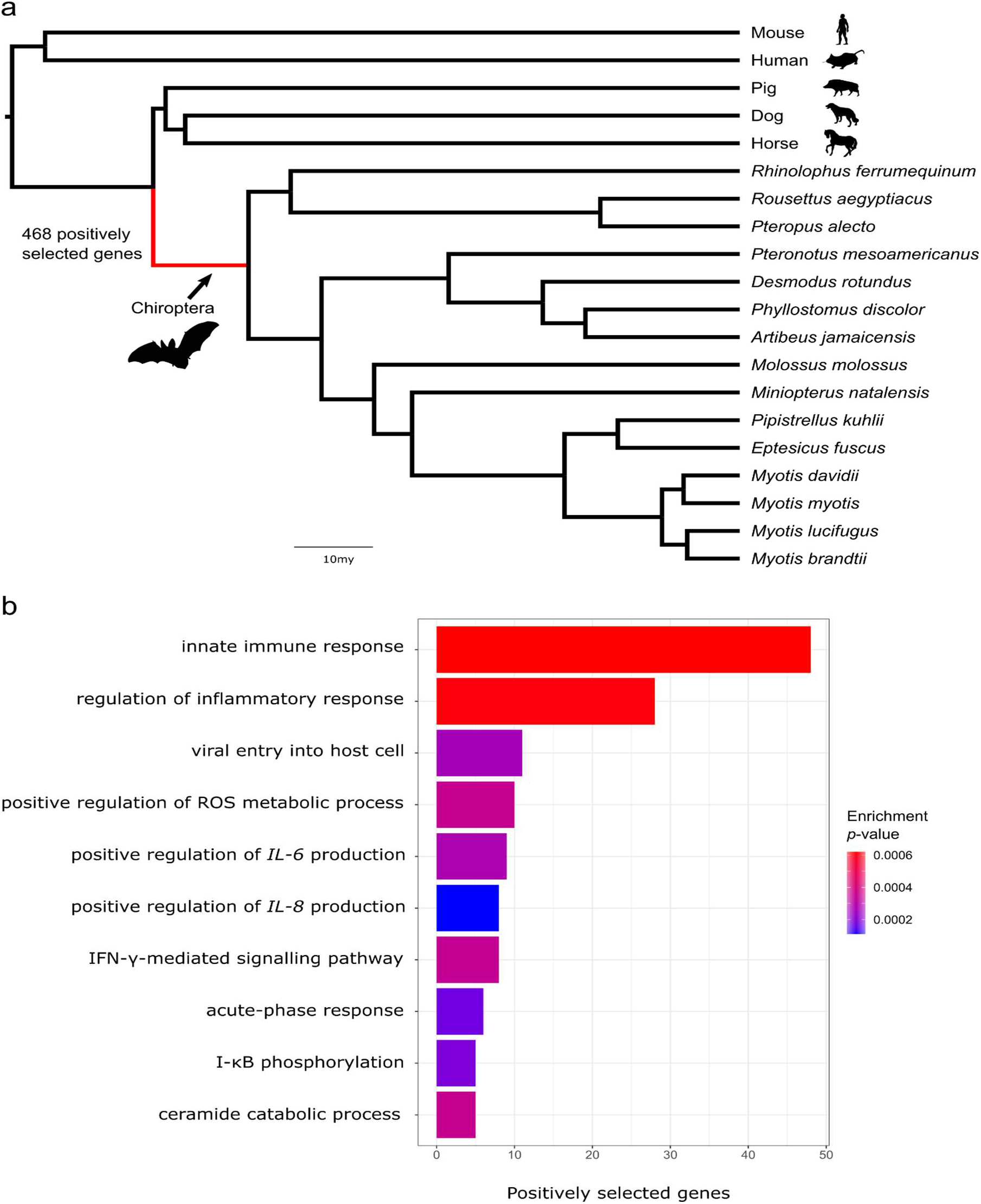
Positive selection scan on the bat ancestral branch. a) Maximum likelihood phylogeny based on codon-site partitioned analysis of 3,632 gene alignments, with the bat ancestral branch indicated. b) TopGO hierarchical gene ontology enrichment analysis of the positively selected genes against the background set of tested genes, suggesting strong enrichment of innate immunity genes. The ten most significant GO terms are shown, eight of which are related to innate immunity including genes involved in interleukin regulation and interferon pathways.

### Pathogen sensors, interleukins, and antiviral genes in bats are rapidly evolving

The 468 PSGs on the bat ancestral branch included several genes that encode proteins with pathogen-sensing roles. For example, the TLR-encoding genes *TLR7* and *TLR8*, are included in our set (**Table S10**), and the related *TLR9* (which shows reduced activation in bats^58^) was on the threshold of statistical significance (*p*=0.05). All of these genes have been identified in previous scans for positive selection^16,23^. Another previously identified PSG in our set is the IFN stimulator *STING*^59^. The TLRs and *STING*, as well as the PSG *NLRP3*^17,60^, all play important roles in inflammation and are considered therapeutic targets for inflammatory disease^61–64^. Positive selection in these genes may play a role in dampening downstream responses to pathogens. Interestingly, the location of a bat-specific substitution in codon 358 of *STING* that was linked to dampened IFN activation^59^ was identified in our scan as one of ten positively selected sites (**Table S7**). In addition to these previously identified PSGs, we identified *TLR2* to be under positive selection. Unlike the nucleic-acid sensing *TLRs 7*,*8,* and *9*, *TLR2* recognizes lipoproteins of pathogens such as bacteria and enveloped viruses^65^. *TLR2* signalling also induces the inflammatory cytokine *TNF-α* in infections with viruses like SARS-CoV-2, and blocking *TLR2* protects against pathogenesis caused by the ‘cytokine storm’^66^. We additionally found evidence of positive selection in the key TLR regulator *UNC93B1*^67^, which is regulated by type I IFNs^68^. *UNC93B1* is thought to regulate nucleic-acid sensing TLRs such as *TLR7* and *TLR9* upstream of the process of TLR trafficking from the endoplasmic reticulum to endolysosomes^69,70^.

Another group prominently represented in our PSGs are genes encoding the interleukins, a collection of cytokines with diverse functions in immunity and inflammation^71,72^. In bats, earlier work identified reduced expression of *interleukin-8*^58^, positive selection of *interleukin-1β,* and loss of *interleukin-36γ*^18^, suggesting that rapid evolution of interleukins may have contributed to unique adaptations in bat immunity. We found that five interleukin-related GO categories were significantly enriched for PSGs in bats (**Table S11**), including “interleukin-1β production” (GO:0032611, *p*=0.00075) and “positive regulation of interleukin-6 production” (GO:0032755, *p*=0.00028). In addition, we identified the pleiotropic cytokine-encoding genes *interleukin-6* and *interleukin-15* as PSGs as well as the genes encoding several interleukin-associated receptors (**Table S10**). *Interleukin-6*, which encodes one of the most important cytokines during infection^73^, showed six sites predicted to be positively selected in bats and a high ratio of nonsynonymous to synonymous mutations on the bat MRCA branch (dN/dS=2.72). Inflammatory cytokine-encoding genes such as *interleukin-6* and *interleukin-15* that were under positive selection in the bat MRCA could be additional contributors to dampened inflammation in bats.

Our PSGs also included several antiviral IFN-stimulated genes, which, as noted above, are activated by type I IFNs^41^ and in several cases (including *APOBEC3* and *Mx*) have been shown to be duplicated^18^ and/or under positive selection^25,74^ in bats. For example, we detected positive selection in *PARP9*, whose gene product interacts with those of *DTX3L* and *STAT1* to enhance IFN responsiveness^75,76^. We also found strong evidence of positive selection, affecting at least 18 amino-acid sites, in *IFIT2*, whose product inhibits replication of a broad range of RNA and DNA viruses^77,78^ (**Figure 4**). Ten of the positively selected sites overlap or are physically close to sites known to impact RNA-binding in the TPR4, TPR5, TPR6 and TPR9 motifs^79^, and a bat-specific lysine-to-methionine substitution occurs at codon 255 which is involved in RNA-binding^79^ (**Figure 4**). Positive selection of *IFIT2* in bats may therefore alter, and possibly enhance, expression of numerous antiviral response genes. Taken together, our results provide novel evidence for rapid evolution in the innate immune system of bats (**Figure 5**).

**Figure 4.**
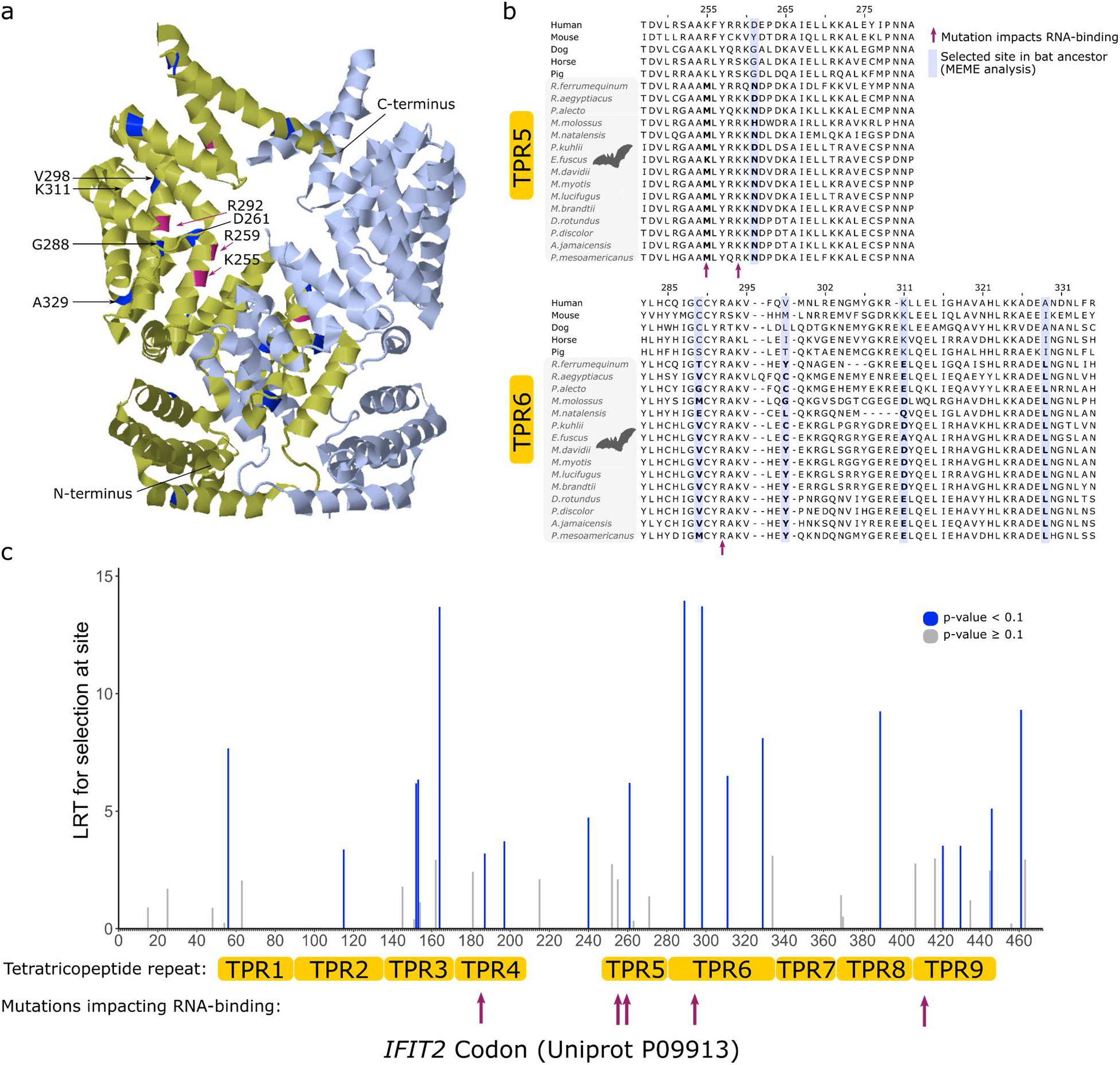
The antiviral gene *IFIT2* is positively selected in RNA-binding regions in bats. a) 3D-structure of the IFIT2 protein (PDB:4G1T) showing sites selected in the bat ancestor (blue) and sites known to be involved in RNA-binding function of the protein (purple). b) Amino acid alignment of the tetratricopeptide repeat (TPR) regions TPR5 and TPR6 (based on UniProt annotation for P09913) showing sites selected in the bat ancestor (blue highlighting and bold) and a fixed bat lysine-to-methionine substitution in codon 255 which is located in a region involved in RNA-binding (indicated with purple arrow). c) MEME^105^ analysis of *IFIT2* codon sites showing 18 sites selected in bats based on a likelihood ratio test.

**Figure 5.**
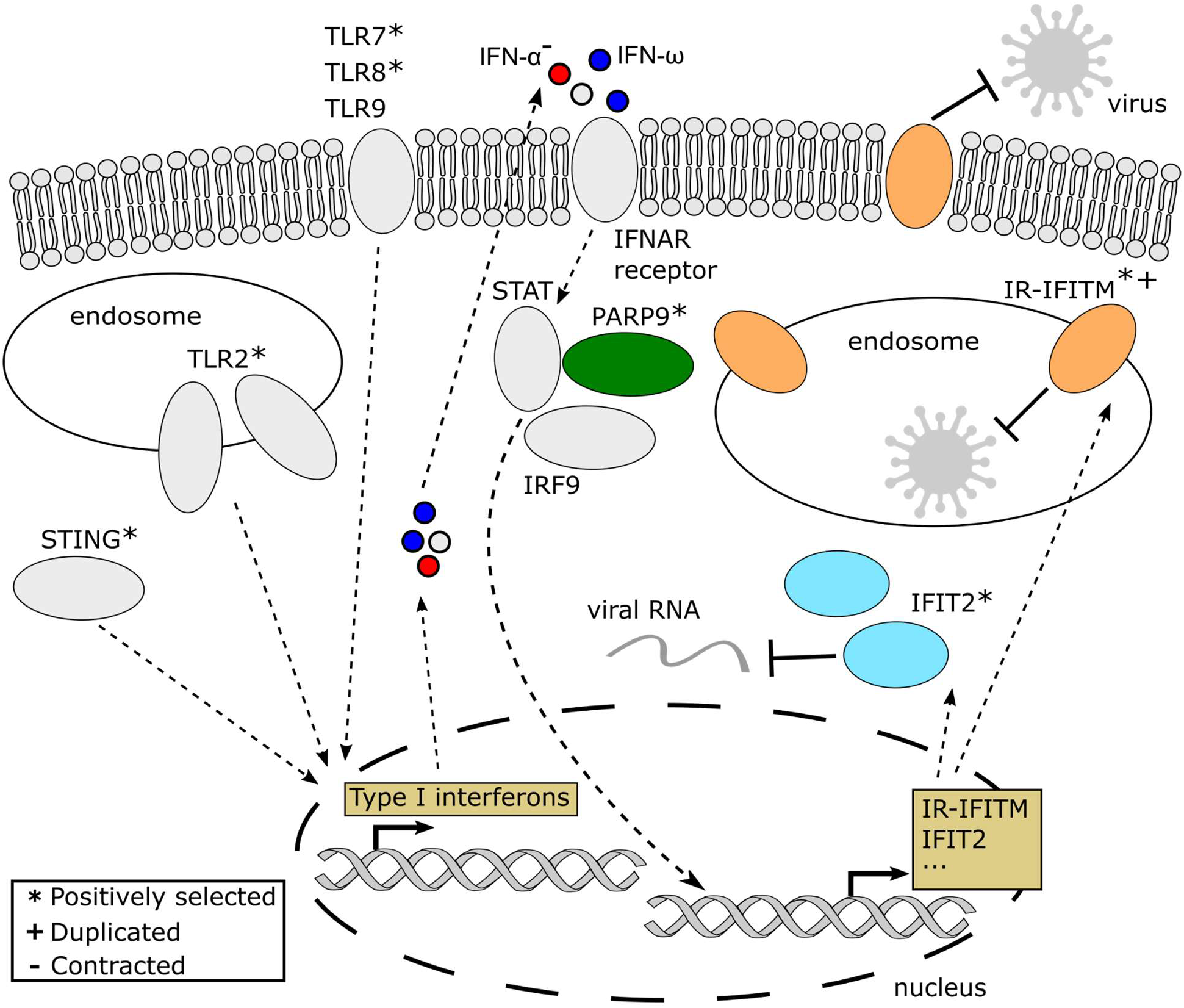
Schema of cellular innate immunity processes associated with genes rapidly evolving in bats. Proteins shown in colour are the most significant innate immunity proteins highlighted in this study. Pathogen sensing pathways involving toll-like receptors (TLRs) and *cGAS*-*STING* can induce expression of type I interferons (IFNs) including IFN-α and IFN-ω. IFN-α genes were lost in the bat ancestor, potentially giving a greater role to IFN-ω. The type I IFNs trigger the induction of IFN-stimulated genes via pathways including *STAT*, which interacts with the positively selected *PARP9*. The IFN-stimulated genes include the positively selected *IFIT2* gene and the immune-related (IR) IFITM genes duplicated in yangochiropteran bats, with both *IFIT2* and IR-IFITM genes playing prominent roles in antiviral defences. The overall schema is based on reviews of type I interferons, PARP9, IFIT and IFITM proteins^78,80^ and studies on IFITM interactions with RNA viruses^81,82^.

### DNA repair genes and tumor suppressors are positively selected in bats

Enhanced DNA repair has been proposed as a mechanism for longevity and cancer resistance in various mammals including bats^27,83^. We identified six DNA repair-related PSGs and 46 PSGs that were “cancer-related” (**Table S10**), meaning they were included in either the Tumor Suppressor Database^84^ or the Catalogue Of Somatic Mutations In Cancer^85^. Notably, cancer-related genes were enriched more than two-fold among PSGs on the bat ancestral branch relative to a set of mammalian branches (Fisher’s exact test: *p*=9e-3, odds ratio=2.2). Among DNA repair genes, we detected evidence of positive selection in the tumor suppressor-encoding *PALB2* and in four DNA polymerase-encoding genes (*POLA1*, *POLD1*, *POLK,* and *POLM*). *PALB2* is a crucial component of the BRCA complex and is required for homologous recombination repair^86–88^. In bats, it shows three sites under selection as well as seven bat-specific coding indels, including a 21-nucleotide deletion in a *RAD51*/*BRCA1*-interacting region (Uniprot annotation: Q86YC2). Despite previous evidence in a long-lived bat^17^, we did not find a signal for selection in *BRCA2*, although it does contain 14 bat-specific indels (**Data S2**). Similarly, we did not identify the DNA repair genes *RAD50* and *KU80* (cf. ref.^17^) or the key tumor suppressor-encoding *TP53* as PSGs, but we did find four sites (codons 35,38,54,97) in *TP53* that are potentially selected in bats as well as a previously described bat-specific indel in the nuclear localization signal domain^17^ (**Figure S7**).

While *TP53* did not appear among our PSGs, we did identify genes encoding two other tumor suppressors that interact with it (**Table 1**): *BIK* and *LATS2*. Both genes showed highly significant signals of selection in bats in our data set but have not been identified in tests of other mammalian branches^89,90^ or in earlier studies in bats^18,20^. *LATS2* is a kinase that modulates the functions of tumor suppressors such as *TP53* as well as canonical growth-related Hippo signaling effectors *YAP*/*TAZ*^91^. We found that *LATS2* is predominantly under negative selection (bat MRCA dN/dS =0.23, outgroup mammals dN/dS=0.11) but nevertheless contains 13 selected nonsynonymous substitutions with evidence of selection in the bat MRCA as well as three bat-specific deletions in its coding region (**Figure S8, Data S2**). Four of the substitutions fall within known functional domains of the protein, with a codon 134 glutamic-acid-to-aspartic-acid substitution in the LATS conserved domain 1 (LCD1) predicted to have an effect on the protein based on SNPeffect^92^. Previous experimental work in mice has shown that the LCD1 domain is critical for tumor suppressor activity of *LATS2*^93^. The second tumor suppressor, *BIK*, belongs to the pro-apoptotic BH3-only family of proteins that are upregulated in response to various stress signals and act as antagonists of pro-survival proteins^94,95^. *BIK* is regulated by *TP53*^96,97^ and contributes to the apoptotic response induced by chemotherapy treatments^94,98^. The rapid molecular evolution of *LATS2* and *BIK* suggests that these genes may contribute to bat adaptations in tumor suppression.

**Table 1.**
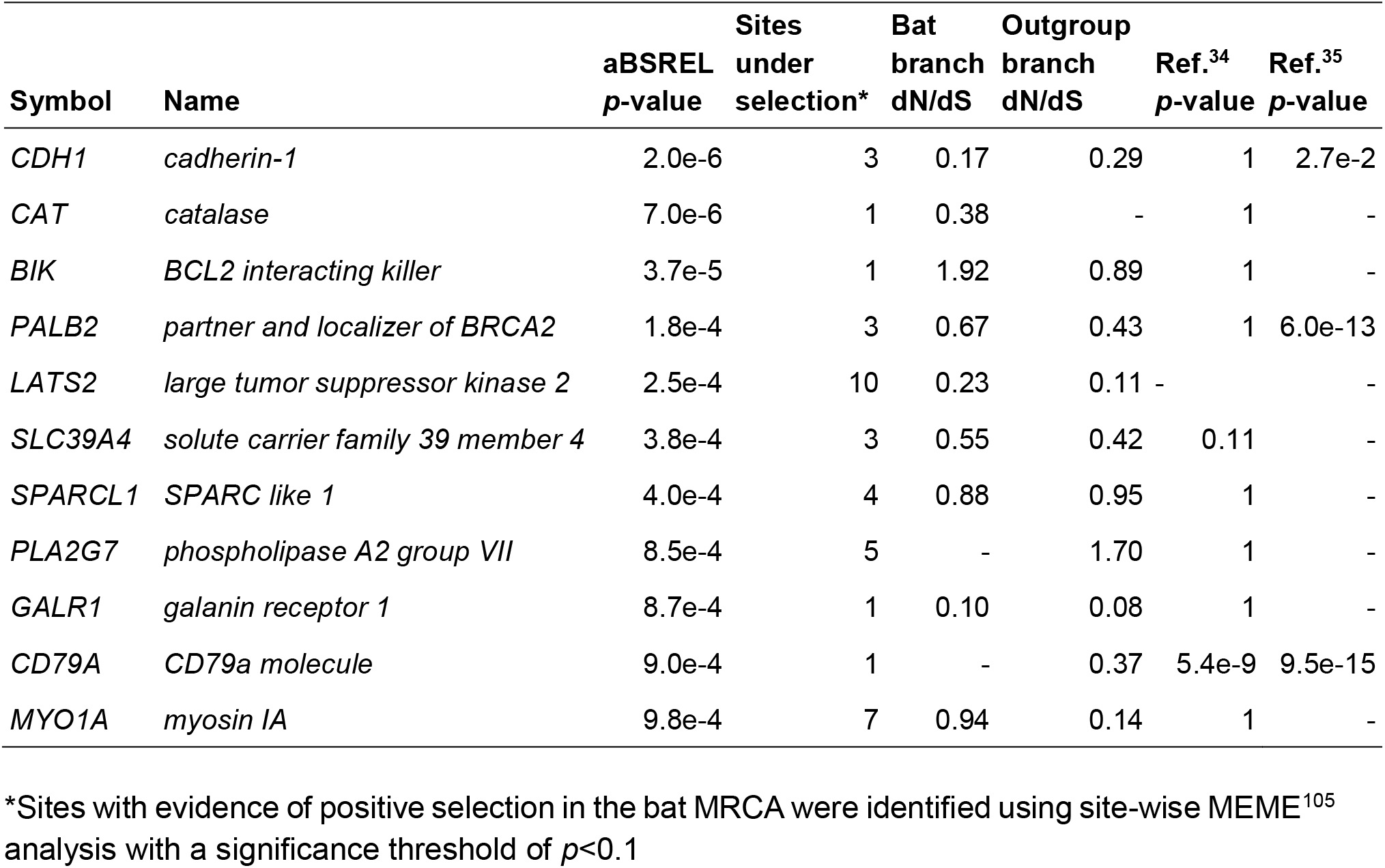
Positively selected genes involved in cancer (*p*<1.0e-3 with at least one site selected in the bat MRCA) that showed the strongest statistical significance. The *p*-values shown (aBSREL *p*-value) are derived from the branch-site likelihood ratio test and adjusted for multiple comparisons across branches but not across genes (see text). For contrast, we also show *p*-values from a similar test (M2a vs. M1a implemented in PAML) applied to a set of mammalian branches in studies by ref.^89^ and ref.^104^. For ref.^89^, nominal *p*-values are shown, with nominal *p*<1.1e−3 corresponding to FDR<0.05; for ref.^104^, adjusted *p*-values are shown, with adjusted *p*<0.05. Missing values are indicated by dashes.

The two cancer-related genes displaying the strongest signals of positive selection, *CDH1* and *CAT* (**Table 1**), also play roles in tumor development. *CDH1* belongs to a family of transmembrane glycoproteins that mediate cell-cell adhesion and regulate cell growth, making them therapeutic targets for preventing tumor progression^99^. *CAT* is an antioxidant enzyme that can protect from ROS-induced stress^100^ and plays a role in *TP53*-mediated ROS regulation in response to DNA damage^101^. Although these functions make *CAT* and *CDH1* intriguing examples of genes that have potentially evolved to enhance bat cancer resistance, previous studies have also found a signal for selection in *CAT*^102,103^ and *CDH1*^102–104^ in other mammals. Experimental validation of the bat-specific mutations detected here will thus be important to demonstrate their potential adaptive value.

## Discussion

High-quality and complete genome sequences are indispensable for revealing patterns of genomic variation within and between species. In this study, we used long-read sequencing to assemble the genomes of the bats *A. jamaicensis* and *P. mesoamericanus* and analysed them together with 13 additional bat genomes to provide insights into unique bat adaptations. We found evidence of genomic adaptation in three key components of the bat innate immune system, including pathogen sensing, type I IFN cytokine signalling, and IFN-stimulated antiviral genes (see schema in **Figure 5**). In particular, the loss of IFN-α genes and the potentially increased reliance of bats on IFN-ω may play a role in their tolerance of viral infections. The expansion of antiviral IR-IFITM genes and *PRDM9* in major bat clades may have further contributed to lineage-specific adaptations. We also found evidence of positive selection in the bat MRCA in 46 cancer-related genes, suggesting a possible link to the unusually low incidence of cancer in bats.

Perhaps our most striking finding, building on earlier comparative genomic studies of the complex type I IFN locus^12,24^, is that eight type I IFNs were lost in the bat MRCA. Notably, we found that bats have lost most, or—in the case of *Pipistrellus kuhlii*, *M. myotis* and *P. mesoamericanus* —all of their IFN-α genes, making their type I IFN locus particularly distinct among mammals. The overall contraction of the type I IFN locus in bats has been hypothesized to allow a smaller number of constitutively expressed IFN-α genes to perform the functions of the 13 IFN-α genes in humans^24^. The constitutive expression and rapid evolution of bat IFNs may also partly result from dampened inflammation caused by evolutionary changes to inflammosome genes such as *AIM2*, *caspase-1* and *IL-1β*^106^, because inflammosomes may negatively regulate type I IFN sensors^107^. Indeed, the regulation of type I IFNs in bats also appears to have evolved adaptively, as positive selection of the key transcription factor *IRF3* in bats was previously shown to enhance antiviral responses via type I IFN activation^108^. However, our results suggest that constitutive expression of IFN-α is not common to all bats, in line with expression analyses in *R. aegyptiacus*^12^. We hypothesize that by relying on the potentially more potent IFN-ω rather than IFN-α, bats may further enhance their antiviral responses. Although further work will be needed to demonstrate a functional shift to IFN-ω, the lack of any IFN-α genes in three bat species strongly suggests a shift has occurred at least in these cases. It is possible that an enhanced antiviral response owing to IFN-ω helps to compensate for an overall dampened inflammatory response in bats. If these properties of IFN-ω can be established, they may open the door to new therapeutic uses of IFN-ω^109,110^.

In addition, our findings suggest that, compared to other parts of the mammalian phylogeny, the lineage leading to bats is enriched for positively selected cancer-related genes. Rapid evolution of DNA repair genes as well as tumor suppressors has been proposed as a mechanism for cancer resistance in other long-lived mammals such as whales^111^. Intriguingly, it was previously hypothesized that selection of cancer resistance components such DNA repair genes in bats resulted from a need to reduce the negative effects of ROS generated as a consequence of flight^17^. This suggestion appears consistent with our findings of positively selected ROS regulators and DNA repair genes. The evolution of cancer resistance in bats may also be associated with adaptations in the bat immune system. There is substantial overlap of genes related to cancer and the immune system^112^, with immune-related genes being known to play a role in cancer surveillance^113^ and tumor suppression^114–116^. Specific examples include immune-related genes that are rapidly evolving in bats. For example, ROS plays an important role in *NLRP3* inflammasome activation^117,118^, *IFIT2* plays a role in apoptosis and cancer progression^119^ and IFNs contribute to anti-tumor activity^120,121^. Positive selection in such genes may be driven by fitness trade-offs between roles in immunity and cancer^112^. For example, inhibition of inflammation can promote longevity^122,123^ and suppress tumor growth^124,125^. Comparative analyses of gene expression across mammals and experimental validation may help resolve the different roles of ROS regulation, DNA repair, inflammation and immunity in cancer resistance. We anticipate that our initial evolutionary findings and the novel genomic resources we have made available (including a genome browser: http://compgen.cshl.edu/bat/) will encourage and facilitate further genomic research in bats, particularly as models that can lead to new strategies for addressing major challenges to human health such as COVID-19 and cancer.

## Acknowledgements

This research was supported, in part, by US National Institutes of Health (NIH) grants P30-CA045508 (to D. Tuveson and colleagues) and R35-GM127070 (to A. Siepel), and by the Simons Center for Quantitative Biology. Computational work was performed with assistance from NIH Grant S10OD028632-01, and the genome browser was set up with help from Ritika Ramani. We thank the CSHL Cancer Center and Director David Tuveson for support and encouragement. We would further like to acknowledge funding support from the CSHL/Northwell Health Affiliation for purchase of the ONT PromethION sequencer used in this study. We also thank Semir Beyaz for helpful comments on the manuscript. Finally, we thank Brock Fenton, Neil Duncan, the staff at the Lamanai Field Research Center, and many other colleagues who assisted with the fieldwork necessary for this study. The content is solely the responsibility of the authors and does not necessarily represent the official views of the NIH or other funding sources.

## Data availability

All raw sequencing data and final genome assemblies have been made available via NCBI (PRJNA751559). Gene annotations can be downloaded or viewed as UCSC genome browser tracks from our laboratory website (http://compgen.cshl.edu/bat/). Supplementary tables and data are available via FigShare (https://doi.org/10.6084/m9.figshare.15223014.v1).

## Materials and methods

### Sample background and acquisition

Fresh liver samples from a single individual of *A. jamaicensis* (AMNH 279493, male) and one *P. mesoamericanus* (AMNH 279536, male) were collected by NBS in April 2017 at the Lamanai Archaeological Reserve in Orange Walk District, Belize (17.75117°N, 88.65446°W). Sampling followed best practices for humane capture, handling, and euthanasia of live mammals outlined by the American Society of Mammalogists^126^. All work was conducted with permission of the Belize Forest Department under permit number WL/2/1/17(19) with IACUC approval from the American Museum of Natural History (AMNHIACUC-20170403) and University of Georgia (A2014 04-016-Y3-A5). Bats were captured in ground-level mist nets and placed in individual cloth bags for transport to the Lamanai Field Research Center. After identification, the bats were euthanized using isoflurane, and the liver was removed immediately after death. Samples were placed in multiple individual 2ml cryotubes and flash-frozen by placement in a liquid nitrogen dry shipper. The cold chain was maintained through shipment to the AMNH, storage in the AMNH Ambrose Monell Cryo Collection, and subsequent sample processing and transfers.

### Genome sequencing and assembly

Approximately 40 mg of liver tissue from each bat was received at CSHL and stored at −80C. The liver samples were crushed with a micro pestle and mixed with 10 ml of TLB buffer and 50μl of RNase A immediately before use. After a one-hour incubation at 37C, 50 μll of Proteinase K was added, followed by incubation at 50C for 3 hours with hourly inversion mixing. After addition of 10ml of a phenol chloroform / isoamyl alcohol mixture, each sample was rocked for 10 minutes and then centrifuged at 4500RPM for 10 minutes. The top aqueous layer was retained and an equal volume of chloroform / isoamyl alcohol was added, and rocking and centrifugation was repeated as above. The top layer was transferred to a fresh tube with 4ml of 5M ammonium acetate. Following the addition of 30ml of ice cold 100% ethanol, the sample was rocked for 10 minutes. The visible DNA was then extracted with a glass pipet and placed in a 1.5ml tube. The sample was washed once with 100% ethanol and centrifuged for 5 minutes at 10,000 RPM. The ethanol was removed and any remaining ethanol was evaporated on a 37C heat block for 10 minutes. The final DNA was resuspended in 10mM Tris HCl pH 8.5 and stored overnight at 4C.

DNA was then sheared to ∼50-75kb using a Diagnode Megarupter following manufacturer’s recommendations. DNA was further enriched for long fragments via the Circulomics small read eliminator XL kit, which iteratively degrades short fragments. DNA was prepared for Nanopore sequencing using the ONT 1D sequencing by ligation kit (SQK-LSK109). Briefly, 1-1.5ug of fragmented DNA was repaired with the NEB FFPE repair kit, followed by end repair and A-tailing with the NEB Ultra II end-prep kit. After an Ampure clean-up step, prepared fragments were ligated to ONT specific adapters via the NEB blunt/TA master mix kit. The library underwent a final clean-up and was loaded onto a PromethION PRO0002 flowcell per manufacturer’s instructions. The flowcells were sequenced with standard parameters for 3 days. Basecalling was performed in real time with Guppy 3.2. Nanopore reads were filtered for minimum length of 10 and minimum 85% accuracy using filtLong (http://github.com/rrwick/Filtlong). Illumina short read libraries were prepared from the same tissue as above with the Illumina TruSeq DNA kit, targeting a 550bp insert size with PCR enrichment.

Libraries were sequenced at the New York Genome Center, on a NovaSeq S4 flowcell in paired end 150bp format to ∼30x genome coverage. Reads were assembled using flye 2.8.3^33^, after evaluation of additional assemblies generated with wtbg2^127^, NextDenovo (https://github.com/Nextomics/NextDenovo) and Shasta^128^ (**Table S12**). Assembly was followed by long-read polishing using minimap 2.17^129^ for alignment and PEPPER 1.0^34^ for polishing. Next, bwa-mem 0.7.17-r1188^130^ was used to align the Illumina short-read data to the long-read polished assembly, and short-read polishing was carried out with POLCA^35^. To compare assembly contiguity, error rates and completeness, assemblies were then assessed using Merqury 1.0^131^, as well as with Benchmarking Sets of Universal Single-Copy Orthologs 4.0.5 (BUSCO)^132^ for mammals (odb9). Additionally, we used python scripts to compute the cumulative sum of contigs length (N(X) length) versus the cumulative sum of N(X)% of the total genome. Further assembly statistics were calculated using BBTools 38.86 (http://sourceforge.net/projects/bbmap/). Duplicated haplotypes in the assemblies were purged using purge_dups 1.2.3^133^.

The assemblies were aligned to human, mouse, and pig assemblies as well as to four bat assemblies (*M. myotis*, *P. discolor*, *D. rotundus,* and *R. ferrumequinum*) using Cactus^134^. Bat-specific small indels (<1kb) were called relative to the human reference from the multiple alignment using a custom python script. Indels were called in alignment blocks with at least 10 bases and seven species aligned including at least 2 non-bat mammals. Only fixed indels present in all bats or all bats but one were retained. Bat-specific insertions and deletions relative to the human reference were distinguished based on the sequence of the pig outgroup.

### Gene annotation

Public RNA-seq data from SRA were downloaded for *A. jamaicensis* and *Pteronotus parnelli* (**Table S13**) and proteins for human (GCF_000001405.39), mouse (GCF_000001635.26), and seven bat species (*Myotis brandtii*, GCF_000412655.1; *M. davidii*,GCF_000327345.1; *M. lucifugus*, GCF_000147115.1; *Phyllostomus discolor*, GCF_004126475.1; *Pteropus alecto*, GCF_000325575.1; *Rhinolophus ferumequinum*, GCF_004115265.1; *R. aegyptiacus*, GCF_001466805.2) were downloaded from RefSeq. RNA-seq reads were aligned to the new reference genomes using HISAT 2.2.0^135^ with the parameters “--no-mixed --no-discordant -- downstream-transcriptome-assembly” and transcripts were assembled using StringTie 2.1.1^136^. To reduce potential loss of transcripts due to limitations of short-read alignment, transcriptomes were also *de novo* assembled with Trinity 2.9.1^137^. PASA 2.4.1^138^ was used to generate the final set of transcripts based on alignment with GMAP^139^ and BLAT^140^ using minimum thresholds of 90% of transcript length aligned at 90% identity. GeMoMa was used to project gene annotations from six bat assemblies^18^ to the genomes of *A. jamaicensis* and *P. mesoamericanus*. Transdecoder was used to predict coding sequences within the assembled transcripts^141^. We used the transcripts and proteins sequences as evidence for the MAKER3 annotation pipeline^142^ with the *ab initio* gene predictors SNAP 2006.07.28^143^ and Augustus 3.3.3^144^. GlimmerHMM^145^ and GeneMark 4.68^146^ were used to generate further *ab initio* gene predictions. Final annotations were generated with Evidence Modeler^147^. To reduce the number of potentially missing genes caused by lack of protein and RNA-seq evidence, we used intact TOGA (https://github.com/hillerlab/TOGA) gene projections from the recently assembled *A. jamaicensis* (GCF_014825515.1) to add gene annotations that did not intersect the Evidence Modeler annotations. The completeness of the final predicted protein set was assessed using BUSCO.

### Repeat analysis

Repeat masking was carried out using an iterative masking and *de novo* repeat detection approach. After masking repeats with Repeatmasker 4.0.9^148^ and the combined RepBase-20181026 and Dfam-3.1 repeat databases, novel repeats were detected in the masked genomes with RepeatModeler 2.0.1. The consensus de novo repeats longer than 100 bp were then concatenated with a vertebrate repeat library including novel bat repeats^18^ and clustered with CD-HIT 4.81^149^. All clustered novel sequences with >80% sequence similarity across >80% of the length of the clustered sequences were excluded^150^. Novel repeats were then aligned to the nt database using BLAST and all repeats matching annotated transcripts were removed. A further BLAST analysis was used to exclude repeats with fewer than 10 alignments to the reference genome from which they were derived. Transposable elements were classified and assigned families using TEclass^151^ and DeepTE^152^. The consensus *de novo* repeats were then concatenated with the vertebrate repeat library and a final masking of the genome was carried out with RepeatMasker using the ‘sensitive’ setting. The recently diverged repeat landscape was analysed using the RepeatMasker script calcDivergenceFromAlign.pl with correction of substitution rates based on the Kimura 2-Parameter model. Transposons were considered recently diverged at 7% divergence from the consensus^18^, which approximates an insertion <30 mya, assuming a mammalian substitution rate of 2.2×10^−9^ ^153^. To allow comparison of the repeat landscape within the Noctilionoidea bat clade, the same repeat masking approach was carried out for several closely related bats (*D. rotundus*, *P. discolor*, S. *hondurensis* and *M. myotis*).

### Mining of endogenous viral elements

To scan for endogenous viral elements that might provide evidence of past infections, we aligned the complete RVDB-prot viral protein database^154^ to our genomes using BLAST with an e-value threshold of 1e-5. Alignments intersecting coding sequence and those shorter than 100aa were excluded. Non-retroviral matches were aligned to the NR protein database using blastx with an e-value threshold of 1e-5 and TaxonKit 0.6.0^155^ was used to retain sequences with a best match to a viral lineage.

### Identification of orthologous gene groups

We analysed bat genes for lineage-specific signals of positive selection and gene duplications based on clustered ortholog groups. Coding sequences for 13 bats and five outgroup mammals (human, mouse, dog, pig, and horse) were downloaded from RefSeq and the Hiller lab (https://bds.mpi-cbg.de/hillerlab/Bat1KPilotProject/). Annotated open reading frames that did not show a nucleotide number that was a multiple of three or that contained internal stop codons were discarded. The longest isoform for each gene was retained. All proteins were clustered with the proteins of our bats using OrthoFinder 2.3.11^156^. A set of single copy orthologs was extracted from the OrthoFinder results, retaining orthologs with at least 12 bat species and three mammalian outgroups. Genes were then aligned using PRANK v.170427^157^ with the species tree provided to guide the alignment. Cancer-associated ortholog clusters were identified based on the databases TSG^84^ and COSMIC^5^.

### Gene family expansion analysis

Alignments of 3,632 single copy genes that were present in all species were concatenated into a single alignment, which was divided into three partitions corresponding to codon positions. A maximum-likelihood phylogeny was inferred using RAxML 8.2.12^158^ rapid bootstrapping under the GTR+G+I model with 100 bootstraps. A dated phylogeny was then generated using MCMCtree in PAML. Node ages were calibrated based on TimeTree^159^ ages for Euarchontoglires, bats, Yangochiroptera and Yinpterochiroptera. Convergence was assessed based on analysis of two replicate runs with tracer^160^. The dated phylogeny was used to calculate gene family expansions and contractions with CAFE 4.21^38^ based on a *p*-value threshold of 0.05. Orthofinder orthogroups were collapsed into 7,405 PANTHER gene families based on BiomaRt^161^ data and 1,448 unassigned Orthogroups. For each of 19,935 orthogroups, a representative gene was selected to obtain a PANTHER assignment. For 2,466 orthogroups, where no human or mouse gene was represented, we selected a gene from a different species for sequence-based annotation using interproscan and eggnog. As CAFE assumes all genes were present in the common ancestor of all analysed species, orthogroups represented in less than two bats and two outgroup mammals were excluded. Final copy numbers and genomic coordinates for IFITM (**Table S14**) and type I IFN (**Table S15**) genes were visualised using genoPlotR^162^. For the analysis of the *PRDM9* expansion in phyllostomids, recently generated short-read assemblies of Sturnira hondurensis (GCA_014824575.2) and A. jamaicensis (GCA_014825515.1) were included in the phylogenetic analysis (**Figure S6**) but not the CAFE analysis. Sequences and RAxML phylogenies are provided for IFITM, IFN, and *PRDM9* orthogroups (**Data S3**).

### Positive selection analysis

We aimed to determine whether each gene was positively selected in one or more of three groups: the bat MRCA, *P. mesoamericanus* and *A. jamaicensis*. We also detected positive selection using the adaptive branch-site random effects model (aBSREL) method implemented in HyPhy 2.5.12^55^. A likelihood ratio test was used to determine whether a lineage-specific group of codons in the alignment is experiencing significant positive selection. Unlike the branch-site tests implemented in PAML^163^, aBSREL allows rate variation among the background branches, which can reduce false positive errors^55^. In addition, we also used PAML 4.9 to apply Test 2, a branch-site test of positive selection. This test compares the alternative model where some branches are under positive selection and thus exhibit sites with ω>1 with the corresponding null model where omega is fixed as 1. We computed *p*-values according to a χ^2^ distribution with one degree of freedom. The *p*-values calculated by PAML and HyPhy were adjusted for multiple testing of three branches per gene using the Benjamini-Hochberg method for controlling false discovery rate (FDR) implemented in base R. We also corrected for multiple testing of all genes using FDR (**Table S10**). We used the maximum-likelihood species tree inferred using 3,048 orthologs (see **Methods** section ‘Gene family expansion analysis’) to provide the topology for the positive selection scans. In this tree, bats are the sister group to Fereuungulata (Cetartiodactyla, Perissodactyla, Carnivora, and Pholidota)^18^. Relationships within Fereungulata remain challenging to resolve^18,164^ and here we followed the TimeTree^159^ topology that places Perissodactyla+Carnivora as sister to Cetartiodactyla. For comparison with a previous scan for positive selection on all branches of a mammalian phylogeny (including human, chimpanzee, macaque, mouse, rat, and dog), we used the set of 8,594 genes that were tested for positive selection using PAML in both this study and in ref^89^. MEME^105^ analysis was used to identify sites potentially under selection in the bat MRCA with a significance threshold of *p*<0.1. Alignment of genes positively selected in the bat MRCA are provided in **Data S4**.

### Gene ontology enrichment

GO, Reactome and KEGG annotations for genes were obtained via BiomaRt. A total of 18,452 orthogroups were annotated with at least one feature and 17,840 were assigned GO features. We carried out enrichment analysis on groups of genes that were positively selected or that showed gene family expansions. Enrichment analysis was performed using topGO 2.44^165^, with the elim algorithm and Fisher’s exact test (*p*<0.01). All genes tested for selection were used as the background. The elim algorithm is a conservative approach that processes the GO graph from the bottom up, to eliminate higher order terms that would otherwise appear enriched due to correlation with more specific terms. GO terms with fewer than 10 genes annotated were excluded.

## Supplementary tables

Supplementary tables are available online (https://doi.org/10.6084/m9.figshare.15223014.v1).

**Table S1**. GenBank assemblies for the taxonomic order Chiroptera (bats). List generated on 4th April 2021.

**Table S2**. Comparison of assembly statistics of the unscaffolded genomes of *Artibeus jamaicensis* and *Pteronotus mesoamericanus* with five scaffolded bat genomes assembled using a range of technologies.

**Table S3**. Recently diverged repetitive elements in the genomes of five noctilionoid bats and the related bat *Myotis myotis*. Repeats were detected and Kimura divergence from the consensus was calculated using RepeatMasker with a custom bat repeat database (available from http://compgen.cshl.edu/bat/).

**Table S4**. Non-retroviral endogenous viral elements (EVEs) detected in the genomes of Artibeus jamaicensis and Pteronotus mesoamericanus. EVEs were identified based on tblastn alignments of the reference viral database (https://rvdb-prot.pasteur.fr/) to the genomes, followed by filtering of sequences <100aa and retroviral sequences. Bat sequences with a best hit to a viral protein or EVE in the NCBI nr protein database using blastp were retained.

**Table S5**. Significantly expanded or contracted gene families in the bat lineage detected using CAFE. Ortholog clusters were assigned to PANTHER gene families based on human protein annotations from biomaRt and gene family descriptions were obtained from pantherdb.org. CAFE results for the Yangochiroptera clade and for *Artibeus jamaicensis* and *Pteronotus mesoamericanus* are also shown.

**Table S6**. Fisher’s exact test using a 2-by-2 contingency table for interferon-α and interferon-ω copy number in a set of outgroup mammals and in individual bat species.

**Table S7**. Significantly selected sites in the genes IFITM3, IFIT2 and STING in the bat lineage based on MEME analysis. Codon positions are based on the human reference sequence. Protein mutagenesis and domain information was extracted from uniprot.org.

**Table S8**. Significantly expanded or contracted gene families in the bat subclade Yangochiroptera and in the species *Artibeus jamaicensis* and *Pteronotus mesoamericanus* detected using CAFE. Ortholog clusters were assigned to PANTHER gene families based on human protein annotations from biomaRt and gene family descriptions were obtained from pantherdb.org.

**Table S9**. Domain analysis of all protein sequences in the *PRDM9* ortholog cluster for seventeen bats and four other mammals. Protein sequences were analysed using PfamScan (https://www.ebi.ac.uk/Tools/pfa/pfamscan/). There is no *PRDM9* ortholog present in the dog. Protein IDs for genomes not generated in this study are based on the GenBank annotations and annotations from https://bds.mpi-cbg.de/hillerlab/Bat1KPilotProject/.

**Table S10**. Positively selected single copy orthologs in bats (order Chiroptera). Gene annotations are based on biomaRt data for the human protein, or, when not available, the protein of mouse or another mammal. Published p-values for the same genes were extracted based on publicly available data from eight studies. Branch-site likelihood ratio test *p*-values are shown for aBSREL and PAML M2a tests for the bat lineage as well as for *Artibeus jamaicensis* (Ajamaicensis) and *Pteronotus mesoamericanus* (Pmesoericanus). Uncorrected *p*-values are shown as well as *p*-values adjusted with FDR for multiple tested branches and for multiple tested branches and genes.

**Table S11**. TopGO gene ontology enrichment analysis of positively selected genes in the bat ancestor. The ‘elim’ algorithm was used and GO enrichments with *p*<0.01 and >=10 annotated genes were retained for further analysis.

**Table S12**. Comparison of assembly and polishing statistics using different approaches. Polished assemblies were assessed using BUSCO v4 (Mammalia odb9) in genome mode and Merqury v1.3. RNAseq reads were aligned using hisat2.

**Table S13**. SRA accession numbers for RNAseq data used in the annotation of *Artibeus jamaicensis* and *Pteronotus mesoamericanus*.

**Table S14**. Genomic coordinates of the genes at the IFITM locus in nine bats and five other mammals. Coordinates for genomes not generated in this study are based on GenBank annotations.

**Table S15**. Genomic coordinates of the genes at the type I interferon locus in 10 bats and five other mammals. Coordinates for genomes not generated in this study are based on GenBank annotations.

## Supplementary data

Supplementary data are available online (https://doi.org/10.6084/m9.figshare.15223014.v1).

**Data S1**. Positive selection analysis of orthogroups in bats (order Chiroptera). All orthogroup clusters generated by Orthofinder are shown, including multicopy orthogroups and lineage-specific orthogroups, but only single-copy orthogroups were analysed for positive selection. Gene annotations are based on biomaRt data for the human protein, or, when not available, the protein of mouse or another mammal. Published *p*-values for the same genes were extracted based on publicly available data from eight studies. Branch-site likelihood ratio test p-values are shown for aBSREL and PAML M2a tests for the bat lineage as well as for *Artibeus jamaicensis* (Ajamaicensis) and *Pteronotus mesoamericanus* (Pmesoericanus). Uncorrected p-values are shown as well as *p*-values adjusted with FDR for multiple tested branches and for multiple tested branches and genes.

**Data S2**. Gene-associated microindels detected in five or more bats from a Cactus multiple alignment of six bats with human, mouse and pig. The bats included in the alignment were *D. rotundus*, *A. jamaicensis*, *P. mesoamericanus*, *P. discolor*, *M. myotis* and *R. ferrumequinum*. Only indels occurring within coding sequences, within 1 kilobase of coding sequence or within the core promoter (based on human core promotors downloaded from https://epd.epfl.ch//index.php) were included. Coordinates are based on the human hg38 assembly and coordinates of aligned sequence from *A. jamaicensis* and *P. mesoamericanus* are also provided.

**Data S3**. Coding sequences and RAxML phylogenies for the type I interferon, interferon-induced transmembrane and PRDM9 orthogroups.

**Data S4**. Unfiltered PRANK alignments of all single-copy orthologs where positive selection was detected on the branch leading to the bat most recent common ancestor.

## Supplementary figures

**Figure S1.**
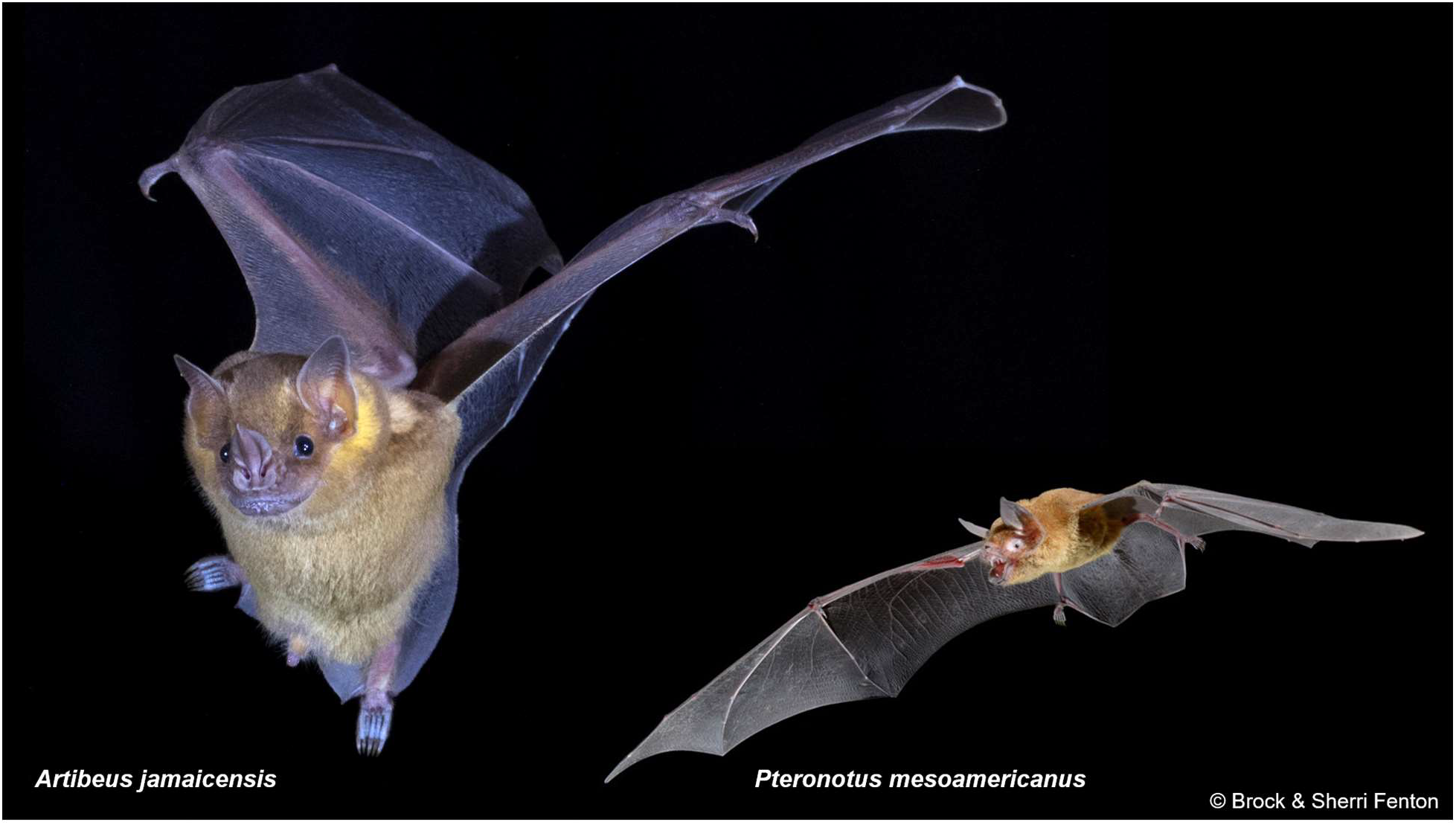
Photographs of the bats *A. jamaicensis* and *P. mesoamericanus* sequenced in this study. Photographs were provided by Brock & Sherri Fenton.

**Figure S2.**
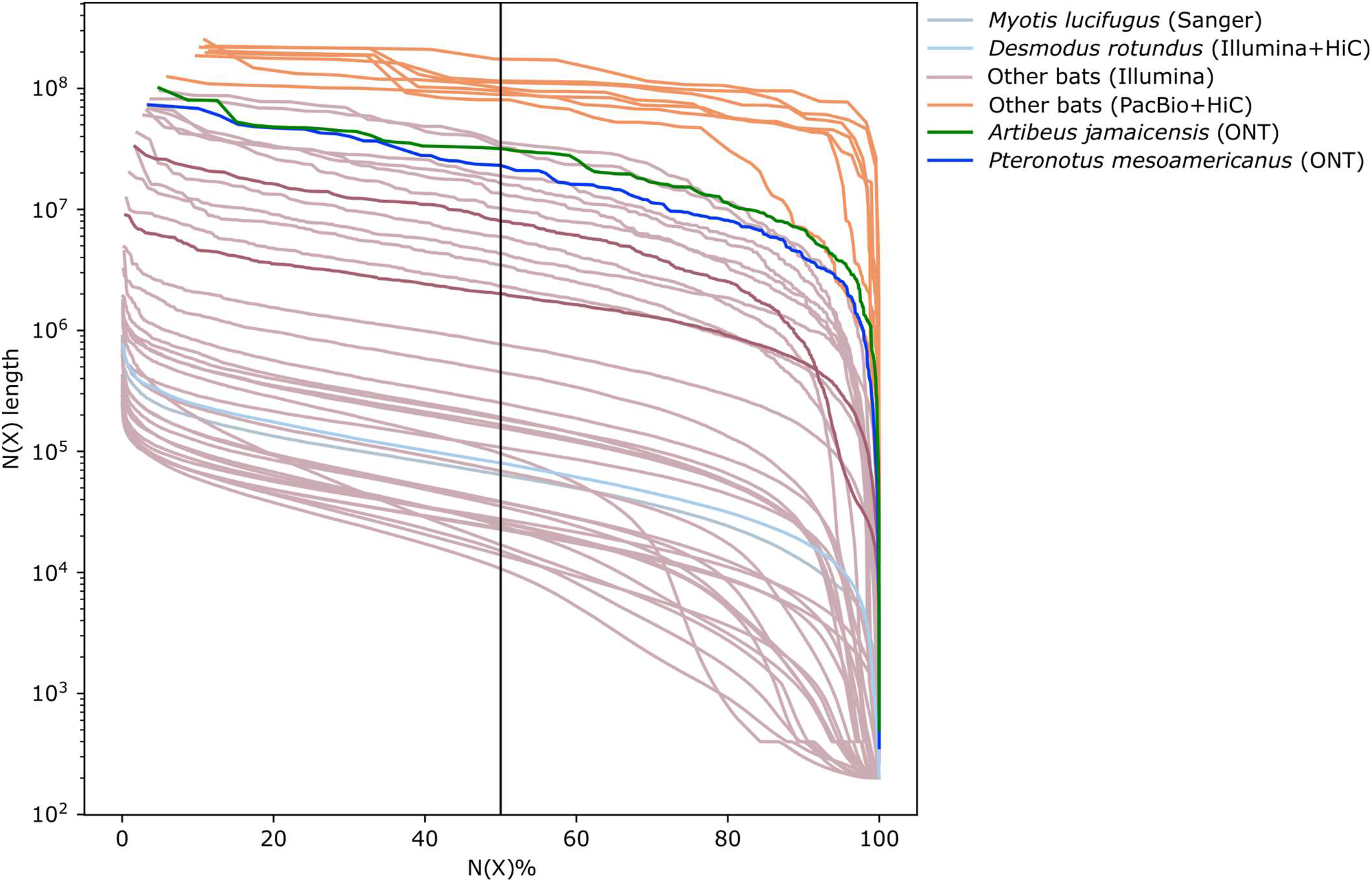
Cumulative sum of the length of the ordered contigs versus the cumulative sum of the total proportion of the genome. The vertical black line indicates the N50 metric. The set of ‘Other bats (PacBio+HiC)’ is based on ref^18^ and ‘Other bats (Illumina)’ are a different set of publicly available bat genomes downloaded from GenBank (see **Table S1** for all GenBank bat assemblies).

**Figure S3.**
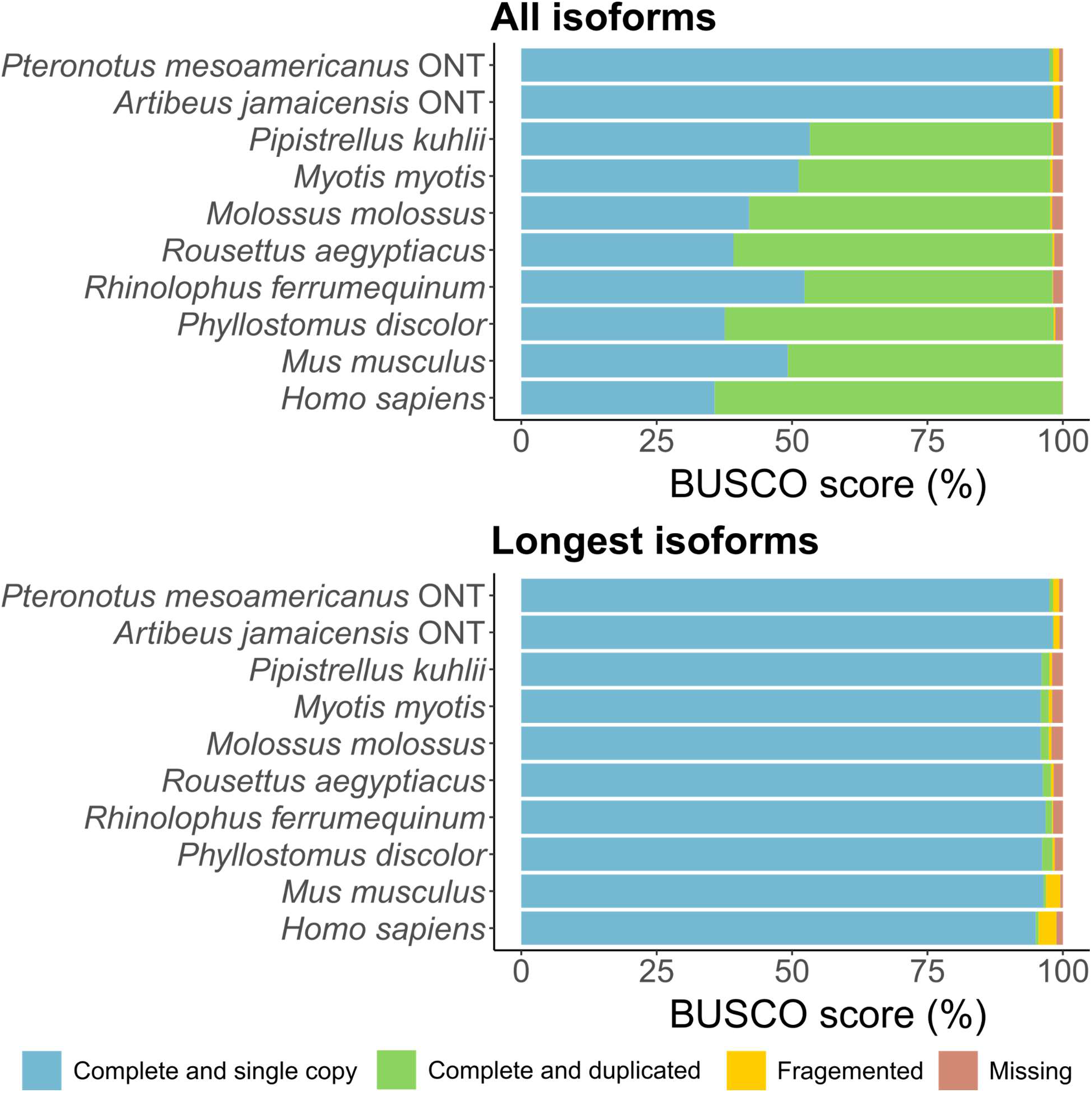
Protein-based BUSCO v4 analysis of eight bats as well as human and mouse. The mammalian BUSCO set (odb9) was used. Proteins for bat species not sequenced in this study are based on annotations downloaded from https://bds.mpi-cbg.de/hillerlab/Bat1KPilotProject/.

**Figure S4.**
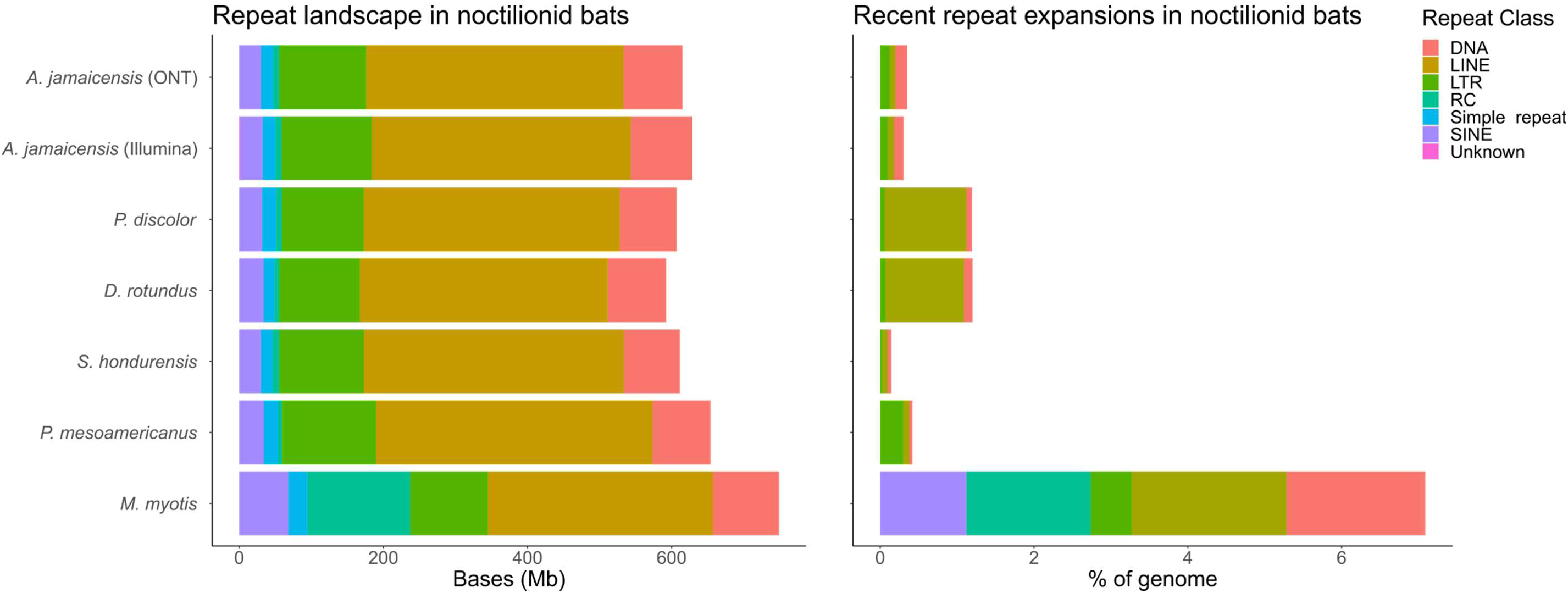
RepeatMasker analysis of repeat classes in noctilionoid genomes. Noctilionoids exhibit a homogeneous landscape of repeat classes that almost completely lacks the rolling circle (RC) repeats prevalent in *M. myotis*. Long interspersed nuclear elements (LINEs) make up the largest repeat class in all bats. Recent repeat expansion make up less than 2% of the genome in all noctilionoids. A list of the recently expanded repeats is provided in **Table S3**.

**Figure S5.**
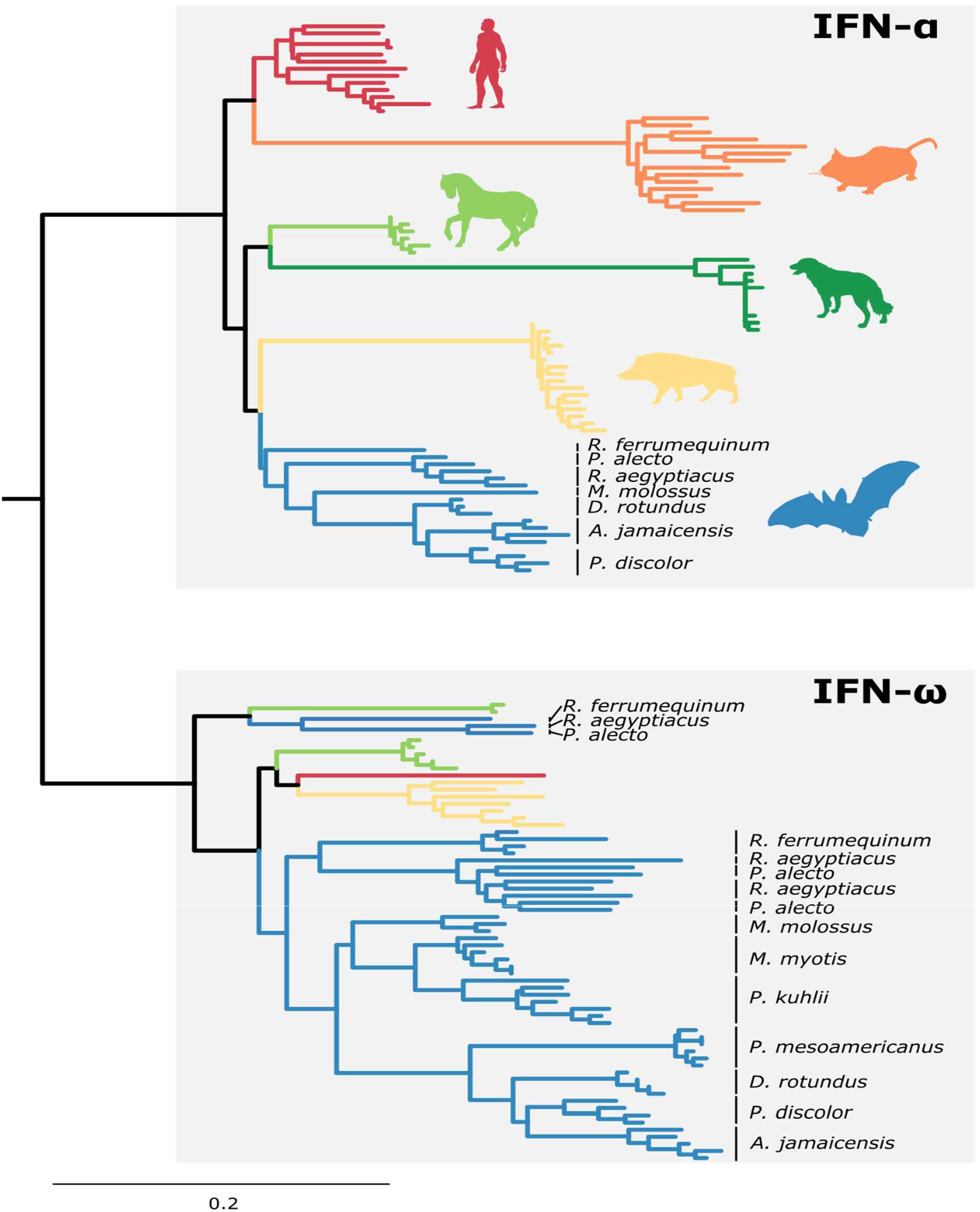
Maximum-likelihood phylogeny of mammalian IFN-α and IFN-ω genes showing the shift in the ratio of IFN-α to IFN-ω copy number in bats (shown in blue) compared to other mammals. The phylogeny was inferred with RAxML under a GTRGAMMA model using alignments partitioned by codon sites. IFN-ω genes are not present in the dog or mouse. Although the topology mostly reflects the expected phylogenetic relationships between species, the tree is intended to show copy number variation in IFN-α and IFN-ω, and it should be noted that internal branches within each ortholog cluster are not robustly supported. The sequence and topology are provided in **Data S3**.

**Figure S6.**
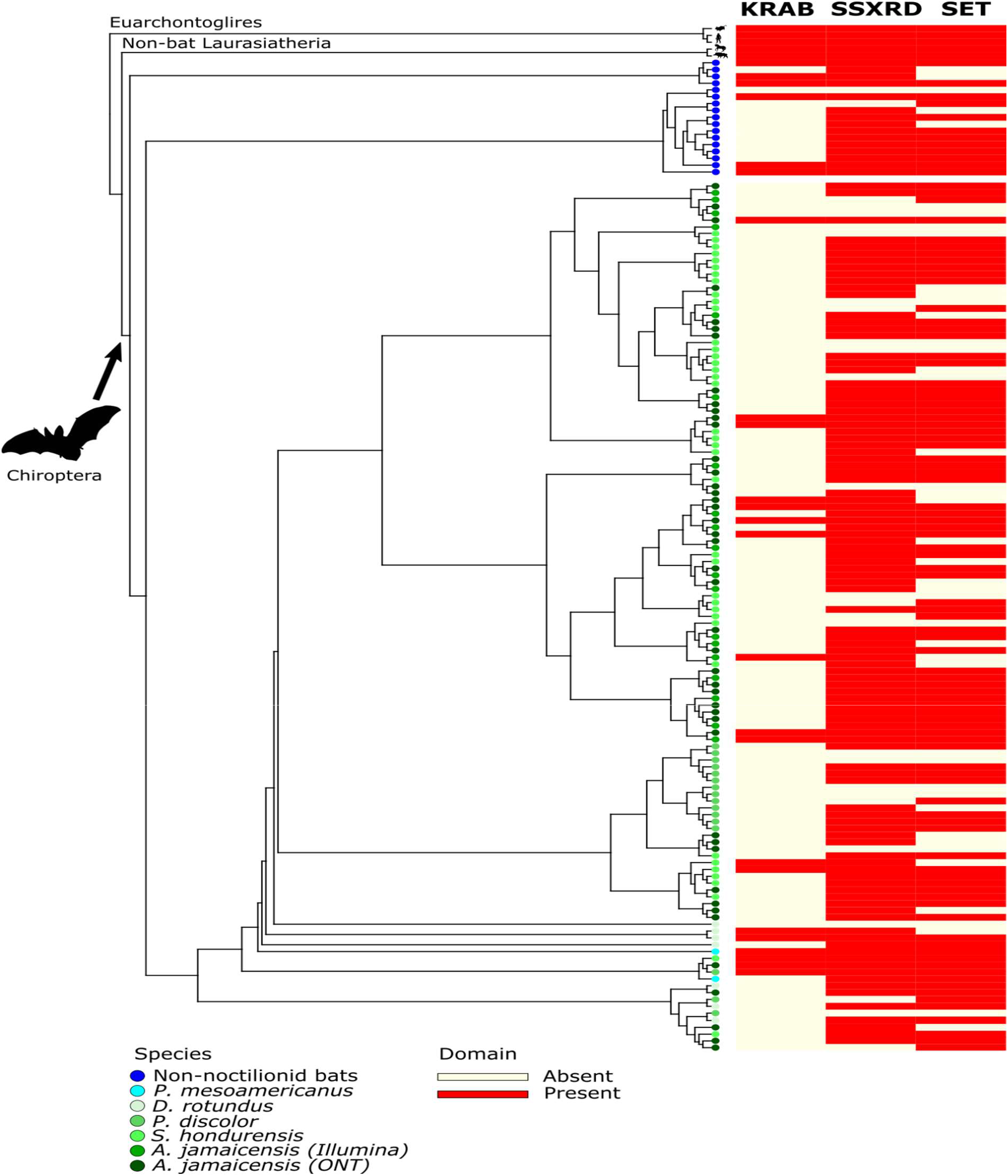
Maximum likelihood phylogeny of the *PRDM9* orthogroup generated using RAxML under the GTRGAMMA model. Presence and absence of the key domains KRAB, SSXRD and SET was determined using pfamscan (https://www.ebi.ac.uk/Tools/pfa/pfamscan/) with default e-value thresholds. The *PRDM9* orthologs underwent an expansion in phyllostomid bats (shown in shades of green). The sequence and topology are provided in **Data S3**.

**Figure S7.**
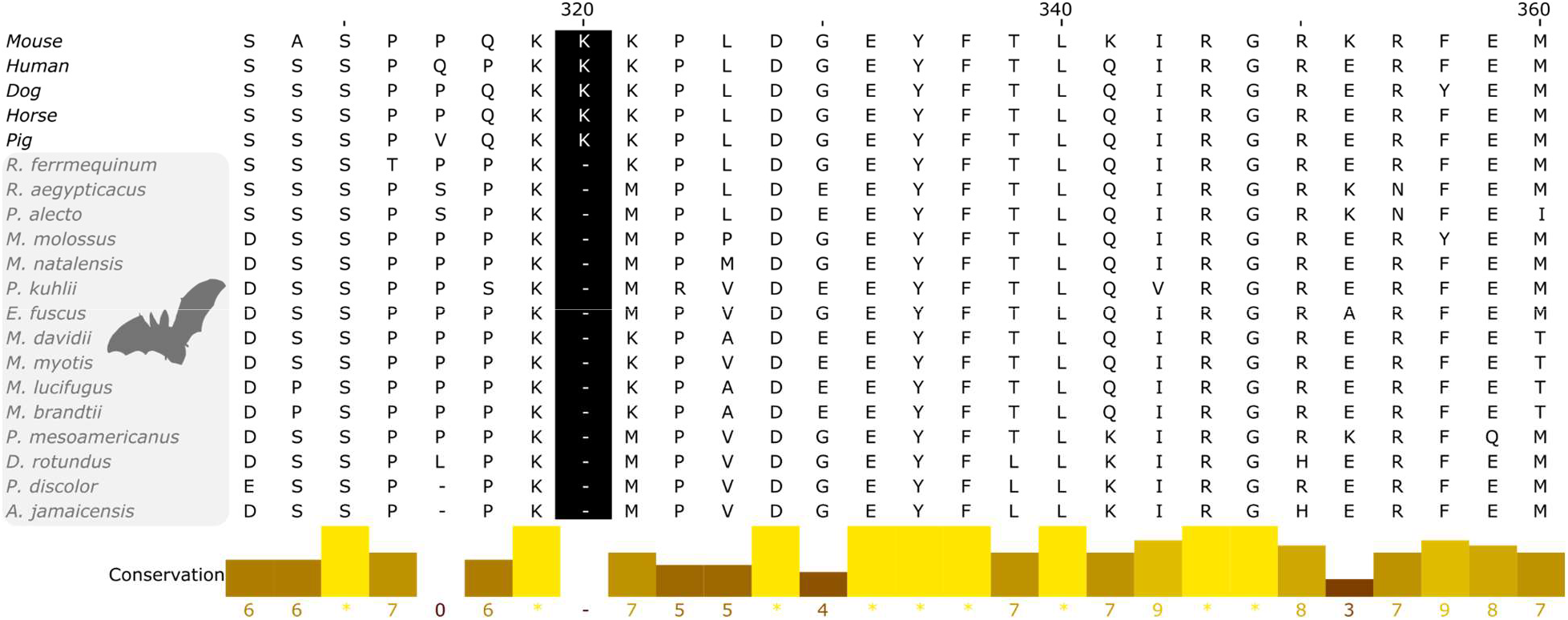
Multiple alignment of *TP53* showing a bat-specific deletion (codon 320) in the nuclear localization signal domain. Amino acid positions are based on the human protein (NP_000537.3). Image was exported from JalView 2.11.1.4.

**Figure S8.**
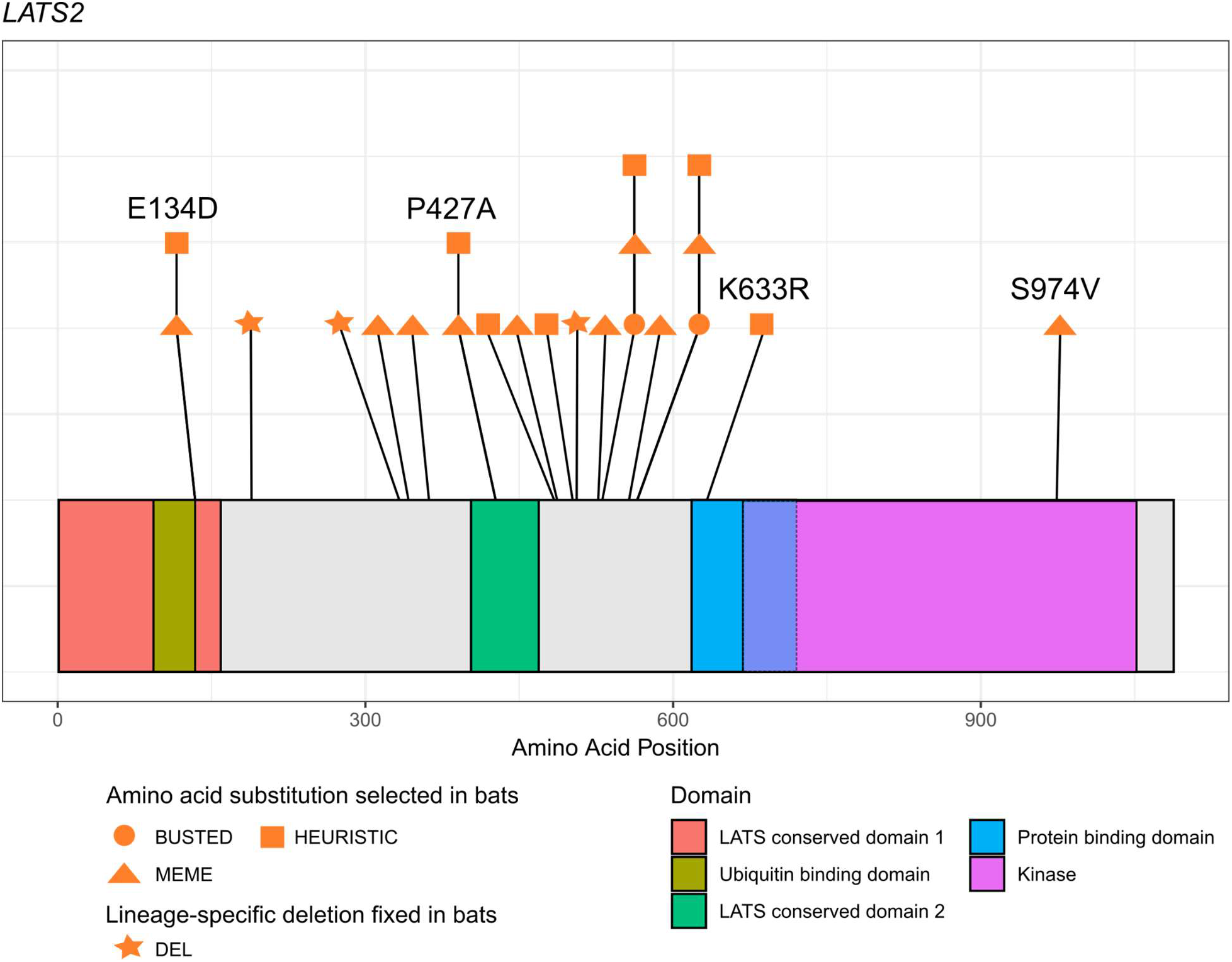
Candidate sites under selection in bats in the *LATS2* gene. Substitutions with evidence of selection were identified using BUSTED and MEME^55^ as well as a simple heuristic (complete bat-specific fixation of a substitution that is not fixed in the outgroup mammals). Bat-specific deletions (DEL) were observed to begin in codons 189, 333, and 506 (coordinates based on human protein XP_016876030.1). Domain coordinates are based on UniProt and ref^93^ (LCD1:1-159, LCD2:403-469, PBD:618-720, KINASE:668-1052, UBA:98-139).

